# AluMine: alignment-free method for the discovery of polymorphic Alu element insertions

**DOI:** 10.1101/588434

**Authors:** Tarmo Puurand, Viktoria Kukuškina, Fanny-Dhelia Pajuste, Maido Remm

## Abstract

**Background:** Recently, alignment-free sequence analysis methods have gained popularity in the field of personal genomics. These methods are based on counting frequencies of short *k*-mer sequences, thus allowing faster and more robust analysis compared to traditional alignment-based methods.

**Results:** We have created a fast alignment-free method, AluMine, to analyze polymorphic insertions of Alu elements in the human genome. We tested the method on 2,241 individuals from the Estonian Genome Project and identified 28,962 potential polymorphic Alu element insertions. Each tested individual had on average 1,574 Alu element insertions that were different from those in the reference genome. In addition, we propose an alignment-free genotyping method that uses the frequency of insertion/deletion-specific 32-mer pairs to call the genotype directly from raw sequencing reads. Using this method, the concordance between the predicted and experimentally observed genotypes was 98.7%. The running time of the discovery pipeline is approximately 2 hours per individual. The genotyping of potential polymorphic insertions takes between 0.4 and 4 hours per individual, depending on the hardware configuration.

**Conclusions:** AluMine provides tools that allow discovery of novel Alu element insertions and/or genotyping of known Alu element insertions from personal genomes within few hours.

## INTRODUCTION

Approximately 45% of the human genome contains repeated sequences. These repeated sequences can be divided into tandem repeats and interspersed repeat elements (segmental duplications and transposable elements). The most abundant transposable element in the human genome is the Alu element. A typical Alu element is an approximately 300 bp long transposable nucleotide sequence. [1–3]. The estimated number of full-length or partial Alu elements in the human genome is 1.1 million [4–7].

The presence or absence of some Alu elements is variable between individual genomes. Members of the Alu AluY and AluS subfamilies actively retrotranspose themselves into new locations, thus generating polymorphic Alu insertions [8–10]. A polymorphic Alu in this context refers to the presence or absence of the entire element and not single nucleotide polymorphisms within the Alu sequence. The insertion rate of Alu elements into new locations is approximately one insertion per 20 births [11,12]. Most of the variation in Alu elements is caused by the insertion of new elements. Deletion of the entire Alu element is possible but occurs much less frequently than the insertion of new elements. Polymorphic Alu insertions disturb the regulation of flanking genes and affect phenotype. They cause changes in the genome that lead to disease [13–15]. Therefore, computational methods that reliably detect polymorphic Alu element insertions from sequencing data are needed.

Several methods for the identification of polymorphic Alu insertions have been developed that include the following: VariationHunter [16,17], Hydra [18], TEA [19], RetroSeq [20], alu-detect [21] and Tangram [22], MELT [23], T-lex2 [24], and STEAK [25]. All these methods are based on the mapping of sequencing reads and the subsequent interpretation of mapping results. The discovery of new insertions is typically based on split locations of a single read and/or the distance between paired reads.

Several databases or datasets that describe polymorphic Alu insertions are available. The oldest resource containing known polymorphic transposable elements is the dbRIP database [26]. It contains insertions detected by comparison of Human Genome Project data with Celera genome data. dbRIP also contains information about somatic Alu insertions that might be related to different diseases. The most comprehensive Alu element dataset is available from the 1000 Genome Project (1000G) [12,27]. A subset of these sequences has been validated by Sanger sequencing [9]. The 1000G dataset is currently the reference set for evaluating the accuracy of structural variant calls generated by other methods. The dbRIP, 1000G, me-scan [28], TEA [19] and HGDP [29] datasets together contain more than 10,000 polymorphic Alu insertions that were collected from hundreds of individuals from different populations.

We have developed a set of novel, alignment-free methods for the rapid discovery of polymorphic Alu insertions from fully sequenced individual genomes. In addition, we provide a method that calls genotypes with previously known insertions directly from raw reads. Evaluation of these methods was performed by computational simulations and PCR product size analysis.

## RESULTS

### Rationale for the alignment-free discovery of Alu insertion sites

We describe a novel method allowing both the discovery of new polymorphic Alu insertions and the detection of known insertions directly from raw reads in next generation sequencing (NGS) data. Two key steps within the discovery method are the a) identification of potential polymorphic Alu insertions present in tested personal genomes but not in the reference genome (REF– discovery) and the b) identification of potential polymorphic Alu elements present in the current reference genome (REF+ discovery) that might be missing in the tested genomes.

All discovery pipelines use a 10 bp consensus sequence from the 5’ end of the Alu (GGCCGGGCGC) with one mismatch that we call Alu signatures. The REF– discovery pipeline identifies all occurrences of Alu signatures in raw sequencing reads from an individual. A 25 bp flanking sequence from the 5’ region is recorded together with the discovered Alu signature sequence (Figure S1 in Additional file 1). Subsequently, the location of these 25 bp sequences in the reference genome is determined using the custom-made software gtester (Kaplinski, unpublished). A new REF– element is reported if the 10 bp sequence in the raw reads is different from the 10 bp sequence in the reference genome.

The REF+ discovery pipeline uses the same Alu element signature to identify all locations in the reference genome where the preceding 5 bp target site duplication motif (TSD) is present 270-350 bp downstream from the signature sequence (see Figure S2 in Additional file 1 for details). Both discovery pipelines generate a pair of 32-mers for each identified Alu element. These 32-mer pairs are used for the subsequent genotyping of the Alu elements in other individuals. Two 32-mers in a pair correspond to two possible alleles with or without the Alu element insertion. All candidate 32-mer pairs are further filtered based on their genotypes in test individuals. The entire discovery process is outlined in Figure 1.

**Figure 1.**
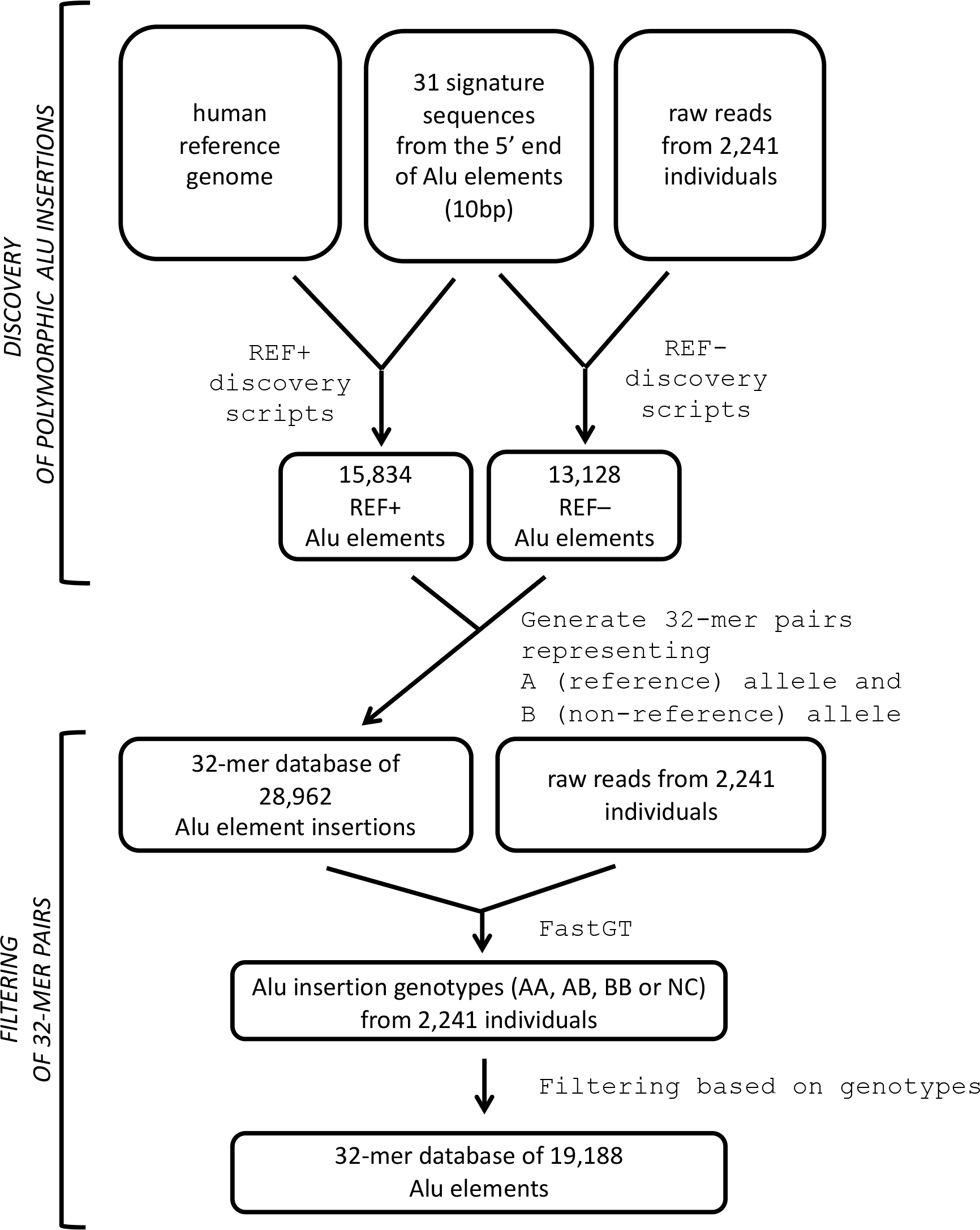
Overview of the discovery methods. Potential polymorphic Alu elements were identified from the raw reads of high-coverage WGS data (REF– Alu elements) and the reference genome (REF+ Alu elements). The candidate Alu elements were filtered using a subset of high-coverage individuals. A final set of 32-mers was used for the fast calling of polymorphic insertions from raw sequencing reads.

The alignment-free genotyping of known Alu elements is based on counting the frequencies of 32-mer pairs specific to Alu element breakpoints using the previously published FastGT software package [30]. The principles of the generation of *k*-mer pairs specific to Alu insertion breakpoints are shown in Figure 2. To detect polymorphic insertions, we use 25 bp from the reference genome immediate to the 5’ end of the potential Alu insertion point and then add either 7 bp from the Alu element or 7 bp from the genomic sequence downstream of the second TSD motif (Figure 2A). The names of two alleles are assigned based on their status in the reference genome; the allele that is present in the reference genome is always called allele A, and the alternative allele is always called allele B (Figure 2B). This allows us to use the same naming convention for alleles and genotypes used by the FastGT package for single nucleotide variants.

**Figure 2.**
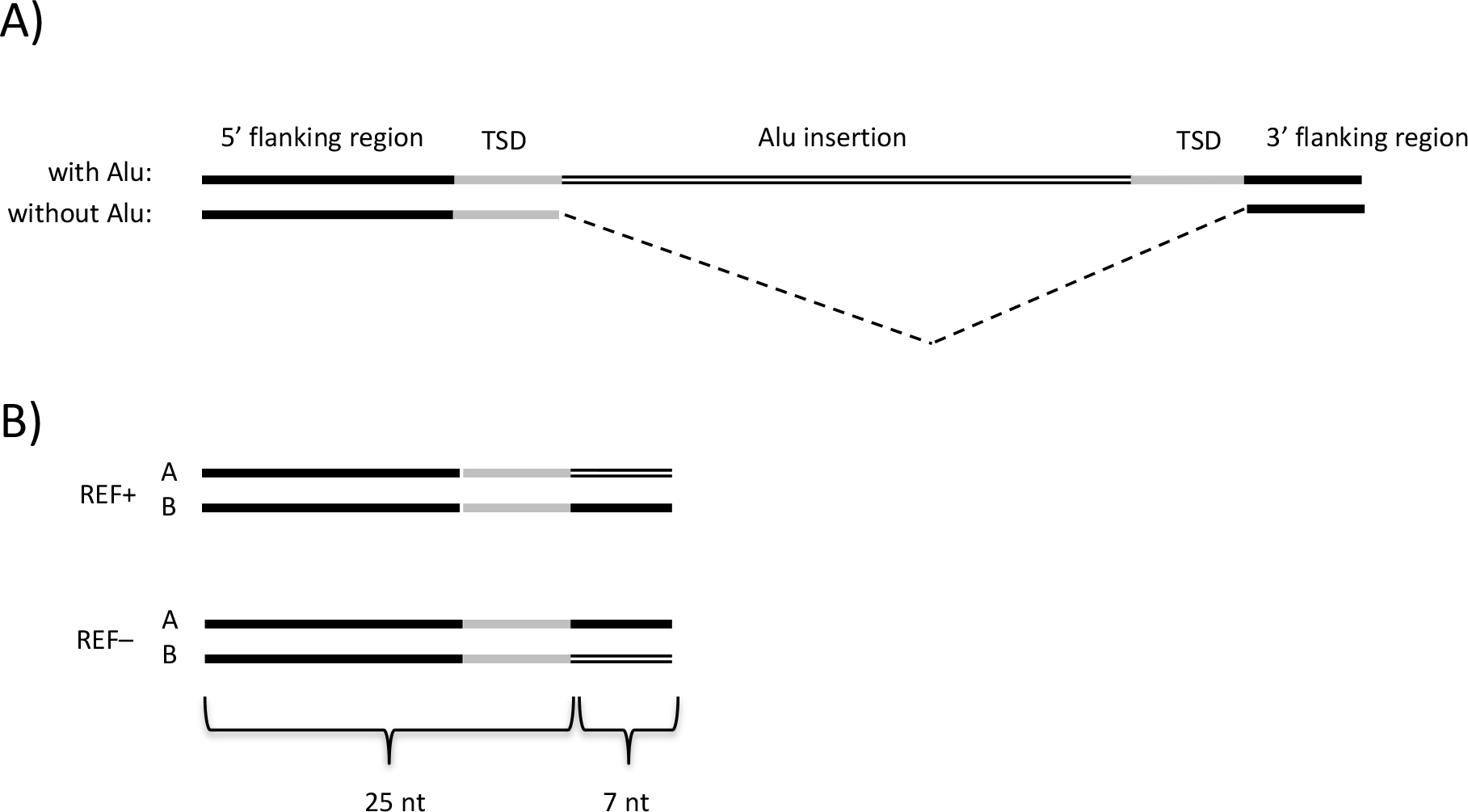
Principle of creating *k*-mer pairs for the calling (genotyping) of polymorphic Alu element insertions. A) Genomic regions with or without an Alu element. B) A pair of 32-mers is created from the insertion breakpoint region covering 25 nucleotides from the 5’-flanking region and 7 nucleotides from either the Alu element or the 3’-flanking region. Allele A always represents the sequence from the reference genome and allele B represents the alternative, non-reference allele.

### Compilation of the list of potential polymorphic Alu elements

To test the applicability of the AluMine method to real data, we performed REF– element discovery using 2,241 high-coverage genomes from the Estonian Genome Project and compiled a set of 32-mer pairs for subsequent genotyping. REF– candidates consist of Alu elements that are present in the raw reads from sequenced individuals but not in the reference genome. We searched the raw reads from test individuals following the principles described above and detected 13,128 REF– Alu elements overall.

REF+ discovery was performed using the human reference genome version 37. We searched for potential REF+ candidates by using the following criteria: the element must have an intact Alu signature sequence, have a TSD at least 5 bp long on both ends of the Alu element, have more than 100 bits similar to known Alu elements, and must not be present in the chimpanzee genome. Our REF+ script detected 267,377 elements with an Alu signature sequence from the human reference genome. However, only 15,834 (5.9%) of these passed all the abovementioned filtering criteria and remained in the set of potential polymorphic elements. The proportion of different signature sequences among the set of REF+ elements is shown in Table S1 in in Additional file 2. All the steps involved in Alu element discovery are summarized in Table 1 together with the number of elements that passed each step.

**Table 1.** Number of REF– and REF+ candidates after different filtering steps

| REF– filtering steps |  |
| --- | --- |
| REF– variations detected in 2,241 individuals | 572,081 |
| REF– candidates that can be located in the reference genome | 379,523 |
| REF– candidates that have unique location in the reference genome | 298,907 |
| REF– candidates after removal of duplicate, closely located and GC-rich k-mers | 13,128 |
| REF– elements that generate reliable genotypes | 9,712 |
| REF+ filtering steps |  |
| Alu signature sequences detected in the reference genome | 267,377 |
| REF+ candidates with 5 bp TSD sequence within 270-350 bp | 110,938 |
| REF+ candidates with BLAST homology | 98,711 |
| REF+ candidates that are not present in chimpanzee genome | 16,434 |
| REF+ candidates after removal of duplicate k-mers | 15,834 |
| REF+ candidates that generate reliable genotypes | 13,396 |

### Simulation tests of the discovery method

We realize that although our discovery methods detected more than 13,000 REF– Alu element insertions, some polymorphic Alu elements remain undiscovered in given individuals. There are two obvious reasons why Alu variants are missed in the REF– discovery step: a) a low depth of coverage in some individuals and b) difficulties with the unique localization of 25-mers in some genomic regions.

The effect of coverage on the discovery rate can be estimated from simulated data. We generated data with 5× to 55× nucleotide-level coverage and analyzed how many REF– elements we would discover from these with our method. The results are shown in Figure 3A. There is an association between the depth of coverage and the discovery rate, which levels out at an approximately 40× depth of coverage.

**Figure 3.**
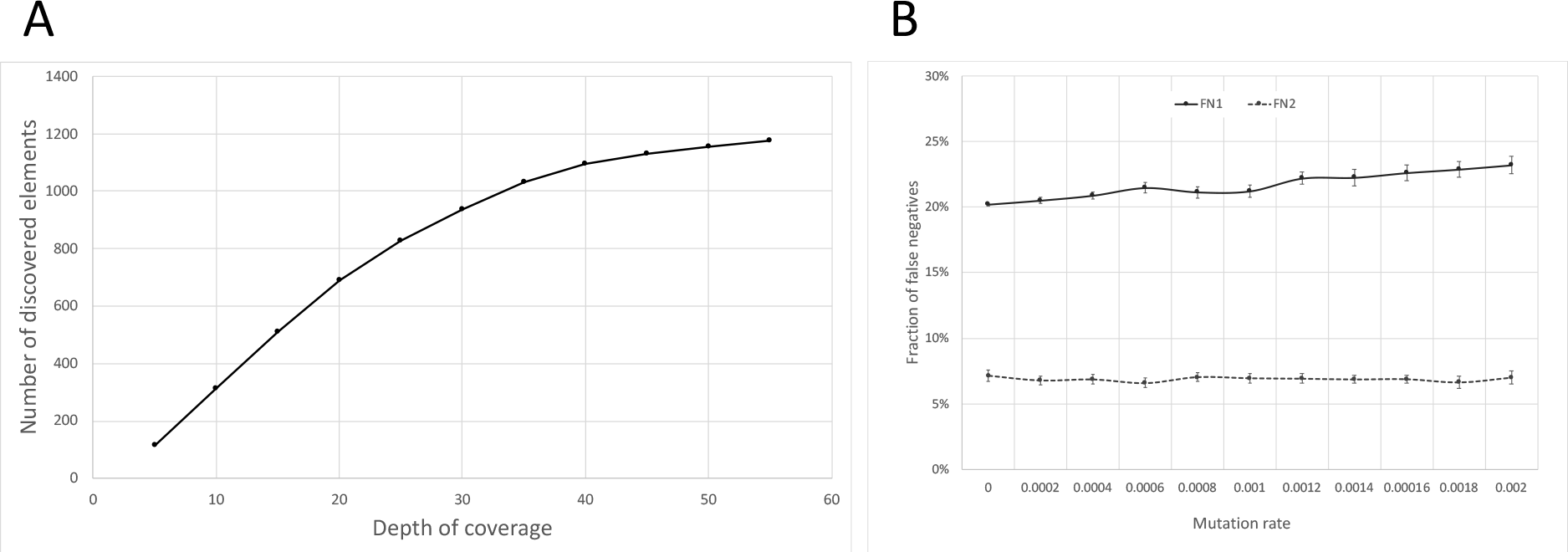
(A) The number of discovered REF– Alu elements in individual NA12877 depending on the depth of coverage. Various depth coverage levels were generated by randomly selecting a subset of reads from the FASTQ file. (B) The frequency of false-negative and false-positive Alu elements found in simulations. FN1 denotes false-negative findings that were undetectable because they are inserted within unsequenced regions of the genome (N-rich regions). FN2 denotes false negatives that could not be detected because they are inserted in nonunique regions of the genome. Error bars indicate 95% confidence intervals from 20 replicates.

Another factor affecting the sensitivity of Alu element discovery is that the repeated structure of the genome sequence prevents the unique localization of discovered Alu elements. The REF– discovery method relies on the unique localization of the 25-mer in front of the Alu signature sequence. We decided to perform a series of simulations with artificial Alu element insertions to determine what fraction of them was discoverable by our REF– discovery method. For this, we inserted 1,000 typical Alu elements into random locations of a diploid genome sequence and generated random sequencing reads from this simulated genome using wgsim software [31]. The simulation was repeated with 10 male and 10 female genomes using different mutation rates. Varying the mutation rate helps to somewhat simulate older and younger Alu element insertions (older Alu elements have accumulated more mutations) and estimate how their detection rate varies accordingly. We observed that 20% to 23% of the elements remain undetected, depending on the mutation rate (Figure 3B). The mutation rate has only a moderate effect on the sensitivity of detection; thus, we assume that the age of the Alu element insertion does not significantly influence the number of detected elements. Additionally, 7% of the inserted elements remained undiscovered because they were inserted into inaccessible (N-rich) regions of the reference genome, and this number is independent of mutation rate.

### Comparison with other Alu discovery methods

When comparing the results of Alu discovery methods, we can compare two aspects. If the same individuals are studied by many methods, we can estimate the overlap between identified elements. Otherwise, we can compare the overall number of detected elements.

We were able to identify the overlap between Alu elements discovered from sample NA12878 within the 1000G pilot project and the 1000G Phase3 project. AluMine discovered 60% (1204) of all elements reported in the 1000G Pilot phase project plus an additional 443 elements (Figure 4). The overlaps between methods are similar for REF+ and REF– elements.

**Figure 4.**
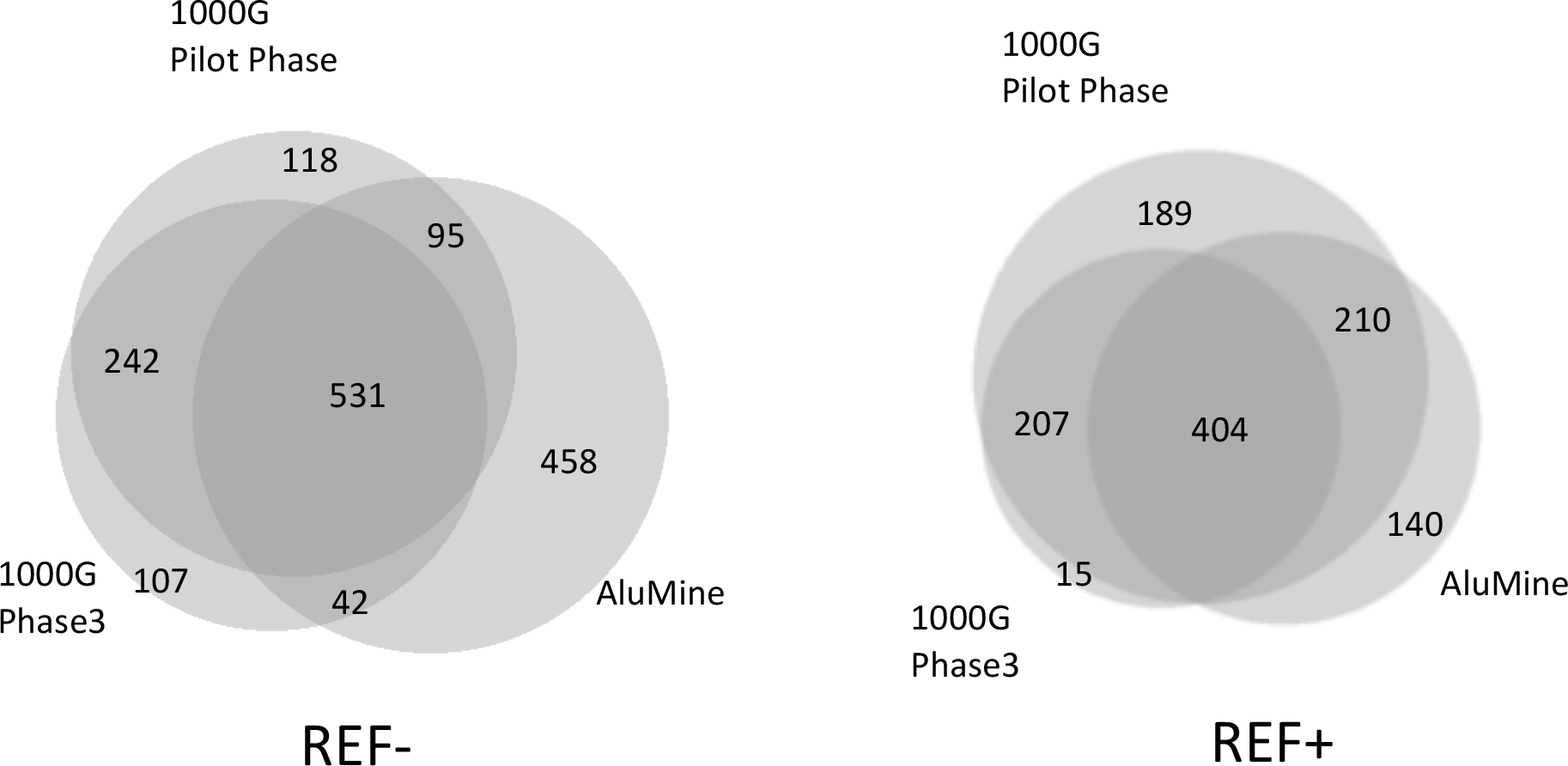
Overlap between REF+ and REF– elements detected by AluMine, the 1000G pilot phase and 1000G Phase 3. The Venn diagram was created with BioVenn software [34].

To examine other methods, we were only able to compare the overall number of discovered REF– elements. AluMine detected 1,116 and 1,127 REF– insertions in the CEPH individuals NA12877 and NA12878 and 1,290 insertions in NA18506. alu-detect discovered on average 1,339 Alu insertions per CEU individual [21]. Hormozdiari et al. detected 1,282 events in the CEU individual NA10851 with 22× coverage and 1,720 events in the YRI individual NA18506 with 40× coverage [16]. TEA detected an average of 791 Alu insertions in each individual genome derived from cancer samples [19]. In genomes from Chinese individuals, Yu et al. discovered 1,111 Alu element insertions on average [32]. Thus, the overall number of detected REF– elements was similar for all methods.

The number of polymorphic REF+ elements (present in the reference genome) has been studied less thoroughly. The number of human-specific REF+ insertions is at least 8,817 [33]. We identified 15,834 potential polymorphic REF+ elements, of which 1,762 were polymorphic in at least one individual in the studied population.

### Frequency of non-reference Alu elements in tested individuals

We scanned 2,241 Estonian individuals with the final filtered set of Alu elements to identify the genotypes of all potential polymorphic Alu insertions in their genomes. All tested individuals had some Alu elements that were different from those in the reference genome. The tested individuals had 741 - 1,323 REF– elements (median 1,045) that were not present in the reference genome and 465 - 651 REF+ Alu elements (median 588) that were present in the reference genome but missing in given individual (Figure 5).

**Figure 5.**
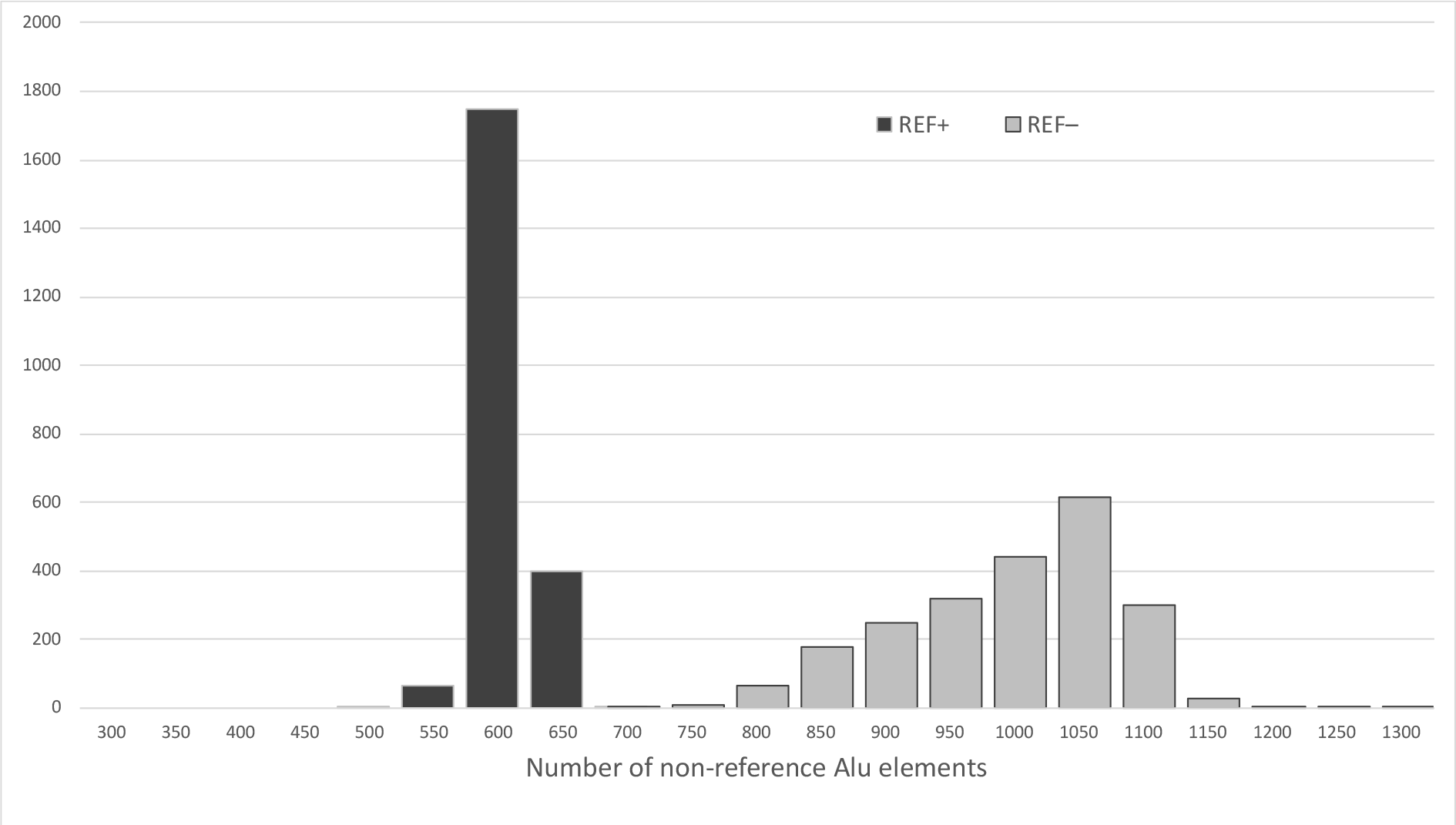
Histogram showing the distribution of the number of non-reference REF– (light) and REF+ (dark) elements discovered per individual genome in 2,241 test individuals from the Estonian Genome Project.

One interesting question that can be addressed from the given data is the cumulative number of REF– elements in a population. We discovered 14,455 REF– Alu elements from 2,241 tested individuals. However, many of these were common within the population. Thus, saturation of the total number of polymorphic elements is expected if sufficient number of individuals are sequenced. The saturation rate of the REF– elements is shown in Figure 6. Obviously, the number of REF– elements was still far from saturation. Each new individual genome sequence still contained 2-3 previously unseen REF– elements.

**Figure 6.**
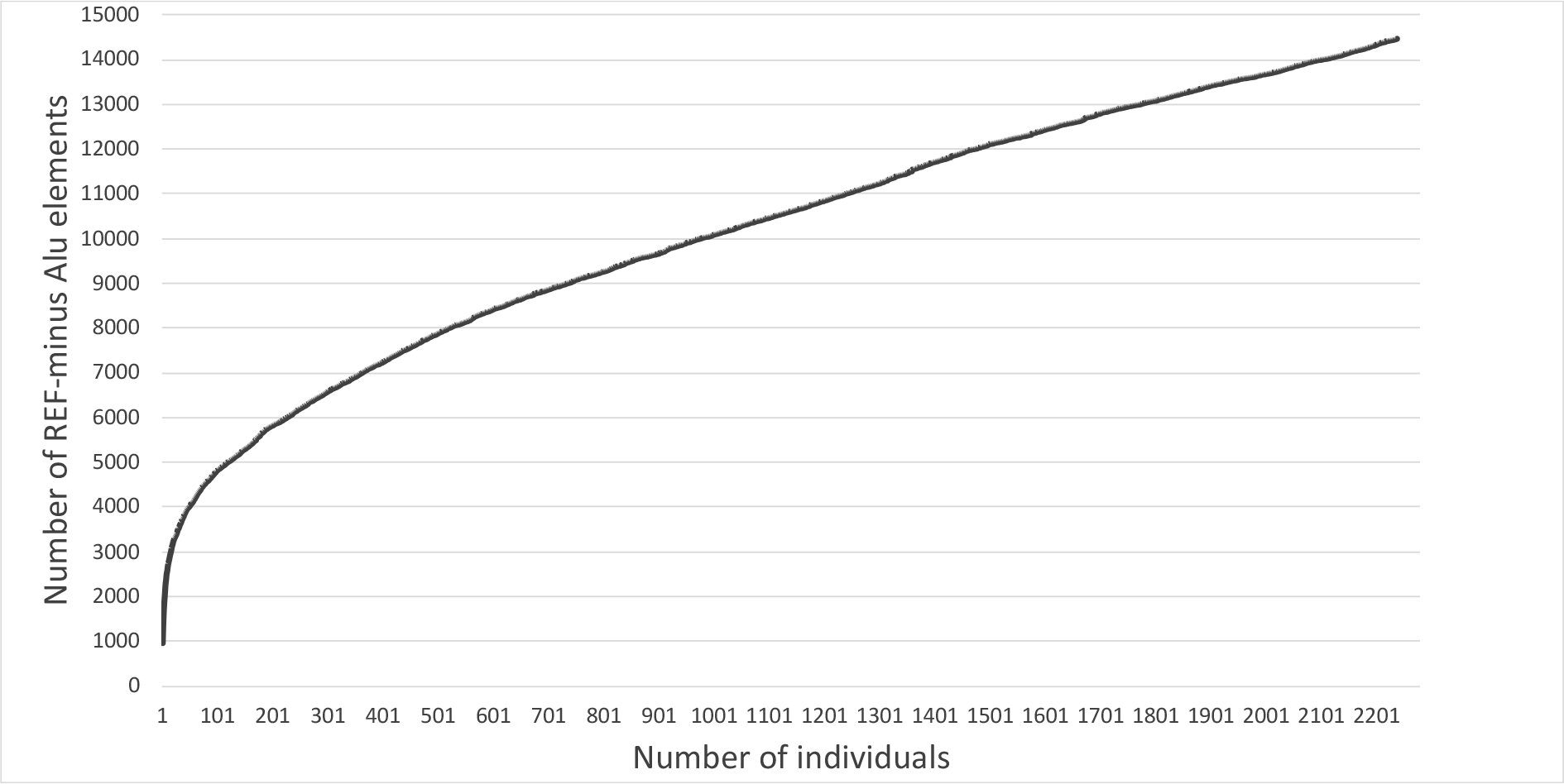
Cumulative frequency of REF– Alu elements discovered from studied individuals.

### Selection of 32-mers for genotyping

In principle, we would like to call the genotypes with discovered Alu elements in other individuals using pairs of specific 32-mers and FastGT genotyping software. Unfortunately, not all discovered Alu elements are suitable for fast genotyping with a pair of short k-mers. Some of them tend to give excessive counts from other regions of the genome, and some might be affected by common Single Nucleotide Variants (SNVs). To select a set of Alu elements that gives reliable genotypes, we filtered the Alu elements based on their genotyping results using data from the same 2,241 individuals that were used for REF– element discovery. To this end, we merged 32-mers of REF– and REF+ Alu elements with a set of SNV-specific 32-mers and determined the genotypes of these markers in test individuals using the FastGT package. SNV-specific *k*-mers are required at this step because Alu elements alone cannot provide reliable estimates of parameter values for the empirical Bayes classifier used in FastGT. Additional filtering and removal of candidate elements was based on several criteria. We removed elements that generated an excessive number of unexpected genotypes (a diploid genotype is expected for autosomes, and a haploid genotype is expected for chrY), elements that deviated from Hardy-Weinberg equilibrium and monomorphic REF– elements. The validation of all tested markers together with their genotype counts is shown in Table S2 in Additional file 2. In the final validated *k*-mer database, we included 9,712 polymorphic REF– elements that passed the validation filters, including 1,762 polymorphic REF+ elements and 11,634 monomorphic REF+ elements. Although 87% of the candidate REF+ elements were monomorphic in the tested individuals, the possibility exists that they are polymorphic in other populations; therefore, we did not remove them from the *k*-mer database.

### Experimental validation of the genotyping method

We decided to validate the alignment-free genotyping of polymorphic Alu elements with a subset of newly discovered Alu elements. The validation was performed experimentally using PCR fragment length polymorphism. We used four different Alu elements (1 REF– and 3 REF+ elements) and determined their genotypes in 61 individuals. The individuals used in this validation did not belong to the training set of 2,241 individuals and were sequenced independently. The electrophoretic gel showing the PCR products of one REF– polymorphism is shown in Figure 7. The results for the three REF+ individuals are shown in Figure 8. The computationally predicted genotypes and experimentally determined genotypes conflicted in only 3 cases; thus, the concordance rate was 98.7%. The 32-mer counts, predicted genotypes and experimental genotypes for each individual are shown in Table S3 in Additional file 2.

**Figure 7.**
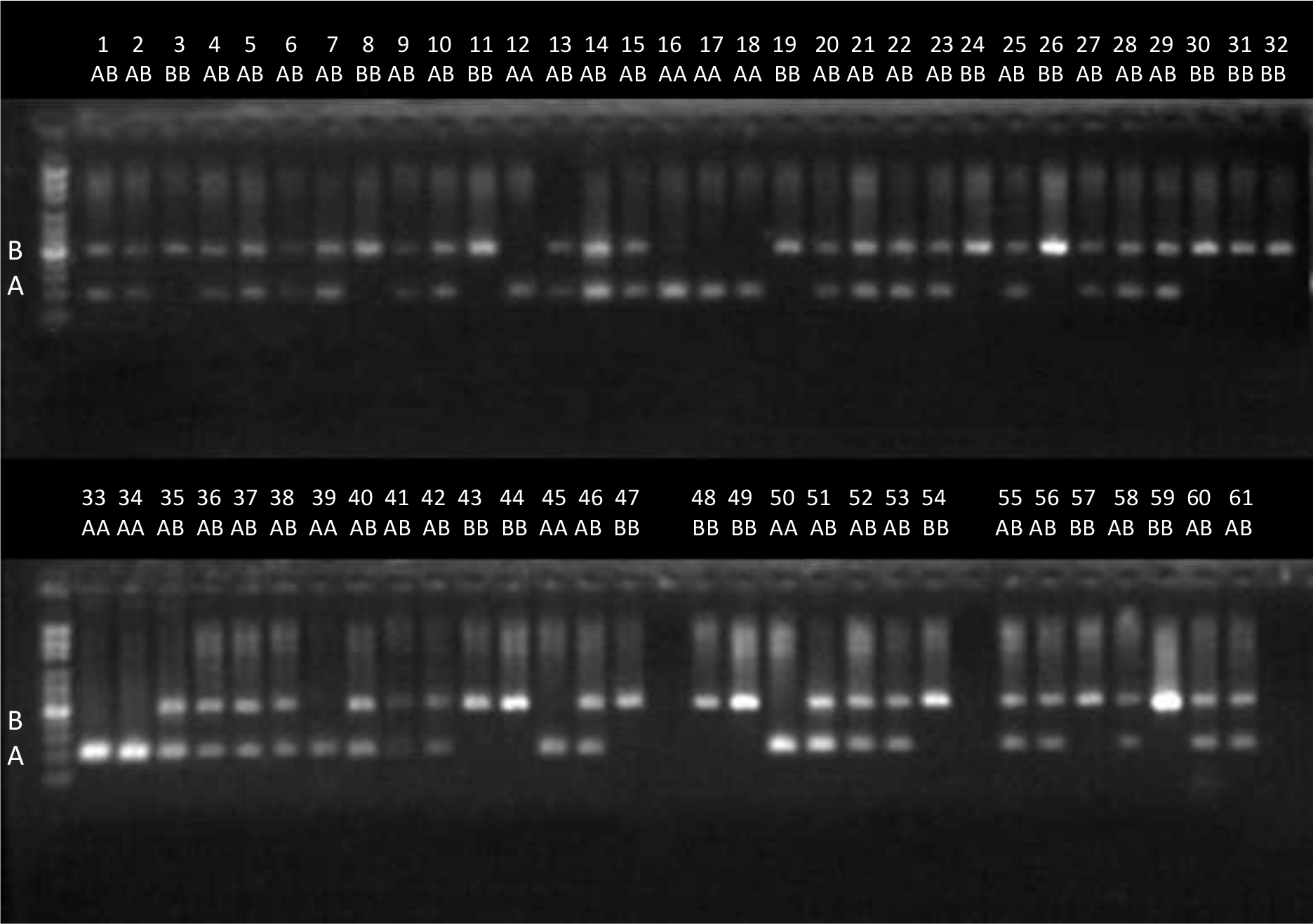
A gel electrophoretic image showing the experimental validation of polymorphic Alu element insertion (REF– elements). One polymorphic Alu element from chr8:42039896 was tested by PCR in DNA from 61 individuals. Lower bands show the absence of an Alu insertion (reference allele A), and upper bands show its presence (alternative allele B).

**Figure 8.**
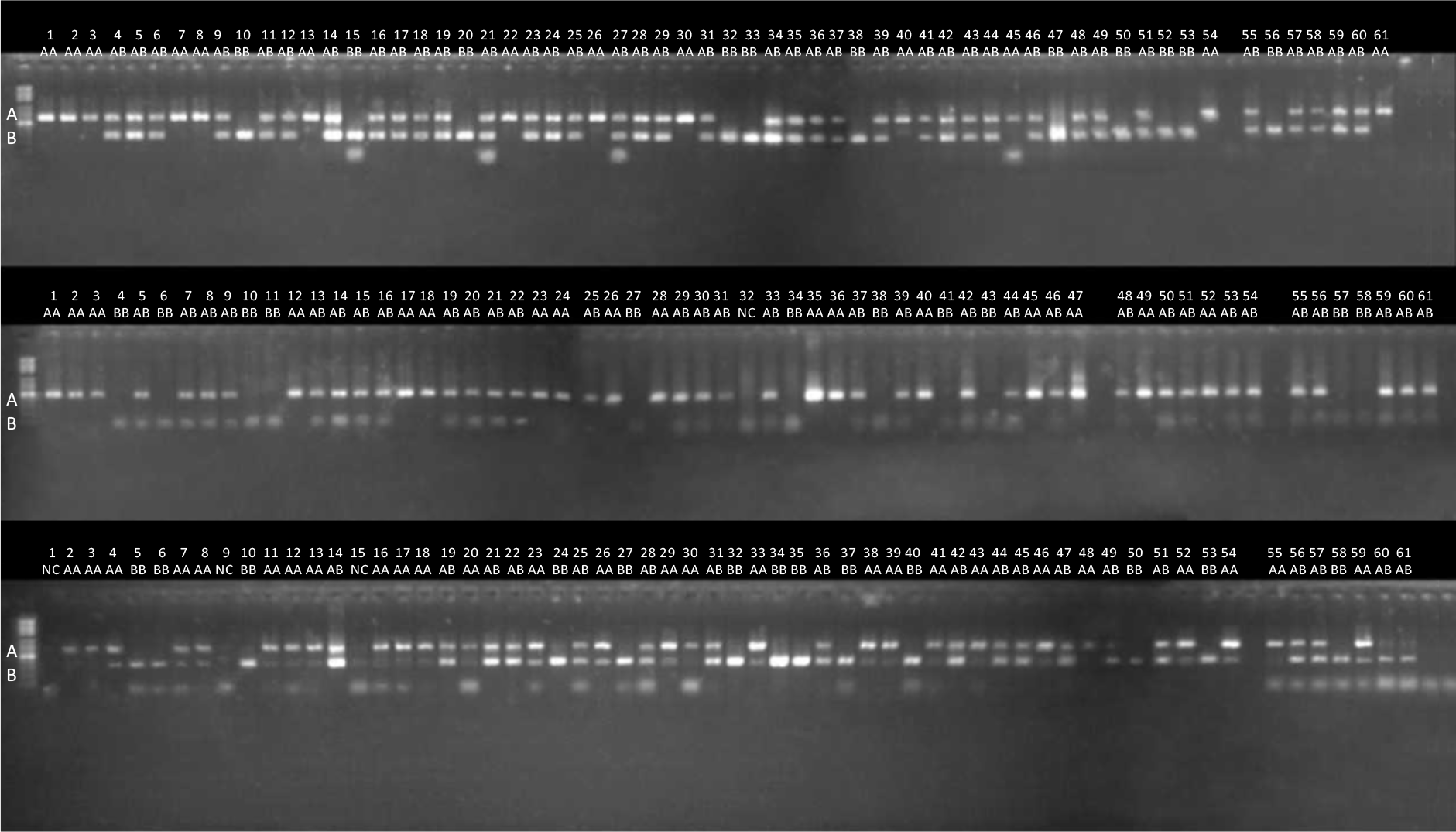
A gel electrophoretic image showing the experimental validation of REF+ polymorphic Alu element insertions. Three locations from chr1:169160349, chr15:69049897 and chr3:95116523 were tested by PCR in DNA from 61 individuals. Upper bands show the presence of an Alu insertion (reference allele A), and lower bands show its absence (alternative allele B).

### Performance

The performance of the AluMine methods can be divided into three parts: the performance of the REF– discovery pipeline, the performance of the REF+ discovery pipeline and the genotyping performance. The REF+ pipeline was run on a server with a 2.27 GHz Intel Xeon CPU X7560 and 512 GB RAM. The REF– scripts and genotyping were run on cluster nodes with a 2.20 GHz Intel Xeon CPU E5-2660 and 64 GB RAM.

The most time-consuming steps in the REF– discovery pipeline are a) searching for Alu signatures from FASTQ files, which takes 2 hours per individual on a single CPU core, and b) finding their locations in the reference genome using gtester software (2 hours for the first individual, 4 minutes for each subsequent individual). The increase in speed for subsequent individuals is due to the large size of the gtester indices (approximately 60 GB). For the first individual, they are read from a hard drive, and for subsequent individuals, the disk cache is used. None of the steps require more than 8 GB of RAM.

The REF+ discovery pipeline contains the following three time-consuming steps: a) a search for 31 different Alu signatures from chromosomes of the reference genome (takes 14 minutes), b) a homology search with all the candidates to confirm that they are Alu elements (2 minutes) and c) a comparison with the chimpanzee genome to exclude fixed Alu elements (4 minutes, 28 GB RAM). All these steps use a single processor. The REF+ discovery pipeline has to be run only once and should not be repeated for each separate individual. Thus, in terms of performance, it occupies only a minor part of the overall analysis.

The genotyping of individuals is performed with the previously published FastGT package [30]. The performance of FastGT was analyzed in the original paper. In optimized conditions (>200 GB RAM available, using FASTQ instead of BAM format, and using solid state drive), it can process one high coverage individual within 30 minutes. However, we used FastGT on cluster nodes with a limited amount of hard drive space and limited RAM. Therefore, in our settings, FastGT acquired sequence data from BAM files through standard input, which limited its performance. In this way, we were able to process one individual in 3-4 CPU hours.

## DISCUSSION

### Parameter choice

A common matter of discussion for alignment-free sequence analysis methods is the optimal length of *k*-mers. In our case, the *k*-mers used for genotyping Alu elements had to be bipartite and contain sufficient sequence from the genome and a couple of nucleotides from the Alu element (Figure 2). The first part of the bipartite *k*-mer must guarantee the unique localization of the *k*-mer in the human genome; the second part must allow distinguishing variants with and without the Alu element at a given location. Both parts must fit into 32 nucleotides because we use the *k*-mer managing software package GenomeTester4, which is able to handle *k*-mers with a maximum length of 32 nucleotides. In the current work, we chose to divide 32-mers into 25 + 7 nucleotides. Our previous work demonstrated that all *k*-mers 22 to 32 nucleotides long should perform equally well to analyze variations in the human genome (Figure 5 in [30]). Thus, we assume that we would obtain a rather similar genotyping result with slightly different splits, such as 22 + 10 or 28 + 4 nucleotides. Using fewer than 4 nucleotides from the Alu element would give too high of a chance to have an identical sequence in the reference genome, and the program would not be able to distinguish variants with and without Alu.

### Comparison with other software

We compared the number of REF– elements discovered by different methods. However, the direct comparison of these numbers to our data is complicated because different populations and individuals were used in different reports. The number of discovered insertions was correlated with the individual ancestry of the subjects: generally, fewer Alu insertions were discovered in CEU individuals than in YRI individuals [12]. Additionally, the depth of coverage had a strong effect on the results, as shown in Figure 3A. All methods, including AluMine, detected approximately 1000 REF-elements per genome. The slight differences were likely due to differences in the depth of coverage and the different origins of the samples used.

Different detection methods have different biases. The premature termination of target primed reverse transcription during the replication of Alu elements can generate truncated Alu element insertions that are missing the 5’ end of the element. It has been estimated that 16.4% of Alu elements are truncated insertions [29]. Furthermore, some Alu element polymorphisms appear through the deletion of existing elements (2%) [9] or mechanisms that do not involve retrotransposition (less than 1%) [29]. Our REF+ method relies on the presence of TSDs, and the REF– method relies on the presence of intact 5’ ends in the Alu. Thus, we would not be able to detect those events, which would explain the majority of the differences between our results and the elements detected in the 1000G pilot phase (Figure 4).

### Future directions

In principle, our discovery method can be used to search for novel Alu elements in any whole-genome sequencing data. Transposable elements are known to occur in genes that are commonly mutated in cancer and to disrupt the expression of target genes [13,19]. Our method allows the discovery of novel Alu elements from sequences from tumors and matched normal blood samples, allowing the study of the somatic insertion of Alu elements in cancer cells and their role in tumorigenesis. The precompiled set of 32-mer pairs allows the genotyping of known Alu element insertions in high-coverage sequencing data. This facilitates the use of Alu elements in genome-wide association studies along with SNVs.

The alignment-free discovery method could also be adapted for the detection of other transposable elements, such as L1 or SVA elements. However, the discovery of these elements is more complicated because SVA elements contain a variable number of (CCCTCT)_n_ repeats in their 5’ end, and L1 elements contain variable number of Gs in front of the GAGGAGCCAA signature sequence.

### CONCLUSIONS

We have created a fast, alignment-free method, AluMine, to analyze polymorphic insertions of Alu elements in the human genome. It consists of two pipelines for the discovery of novel polymorphic insertions directly from raw sequencing reads. One discovery pipeline searches for Alu elements that are present in a given individual but missing from the reference genome (REF– elements), and the other searches for potential polymorphic Alu elements present in the reference genome but missing in some individuals (REF+ elements). We applied the REF– discovery method to 2,241 individuals from the Estonian population and identified 13,128 polymorphic REF– elements overall. We also analyzed the reference genome and identified 15,834 potential polymorphic REF+ elements. Each tested individual had on average 1,574 Alu element insertions (1,045 REF– and 588 REF+ elements) that were different from those in the reference genome.

In addition, we propose an alignment-free genotyping method that uses the frequency of insertion/deletion-specific 32-mer pairs to call the genotype directly from raw sequencing reads. We tested the accuracy of the genotyping method experimentally using a PCR fragment length polymorphism assay. The concordance between the predicted and experimentally observed genotypes was 98.7%.

The running time of the REF– discovery pipeline is approximately 2 hours per individual, and the running time of the REF+ discovery pipeline is 20 minutes. The genotyping of potential polymorphic insertions takes between 0.4 and 4 hours per individual, depending on the hardware configuration.

## METHODS AND DATA

### Data

The reference genome GRCh37.p13 was used for all analyses.

### Discovery of REF– and REF+ elements

The exact details of all discovery pipelines are described in the corresponding scripts (pipeline_ref_plus.sh, pipeline_ref_minus.sh and pipeline_merging_and_filtering.sh) available from GitHub (https://github.com/bioinfo-ut/AluMine).

### PCR protocol

To prepare a 20 µl PCR master mix, we mixed 0.2 µl FIREPol DNA polymerase (Solis BioDyne, Estonia), 0.6 µl of 10 mM DNTP, 0.8 µl of a 20 mM primer mix, 2 µl of 25 mM MgCl2, 2 µl polymerase buffer, and 14.4 µl Milli-Q water. For PCR, Applied Biosystems thermocyclers were used. The PCR was run for 30 cycles using a 1 minute denaturation step at 95°C, a 1 minute annealing step at 55°C and a 1.5 minutes elongation step at 72°C. For gel electrophoresis, a 1.5% agarose gel (0.5 mM TBE + agarose tablets + EtBr) was used. The PCR primer pairs used for the amplification of potential polymorphic regions are shown in Table S4 in Additional file 2.

### Simulated Alu insertions

To simulate polymorphic Alu insertions, we inserted 1000 heterozygous Alu elements into random locations of the diploid reference genome together with a 15 bp target site duplication sequence and a random length polyA sequence (5-80 bp). A male genome (5.98 Gbp) and a female genome (6.07 Gbp) were generated by merging two copies of autosomal chromosomes and the appropriate number of sex chromosomes into a single FASTA file. Simulated sequencing reads were generated using wgSim (version 0.3.1-r13) software from the SAMtools package [31]. The following parameters were used: haplotype_mode = 1, base_error_rate = 0.005, outer_distance_between_the_two_ends = 500, length_of_ reads = 151, cutoff_for_ambiguous_nucleotides=1.0, and number_of_reads = 306,000,000.

## ABBREVIATIONS

1000G: 1000 Genome Project
NGS: Next Generation Sequencing
REF–Alu element: polymorphic Alu element present in at least one personal genome but not in the reference genome
REF+Alu element: polymorphic Alu element present in the reference genome, but missing in at least one personal genome
TSD: Target Site Duplication motif
SNV: Single Nucleotide Variant

## DECLARATIONS

### Acknowledgements

The authors thank Lauris Kaplinski for advice on improving performance of Alu element discovery algorithms and for adapting the *k*-mer counting software FastGT and gtester for this project.

### Authors’ contributions

TP conceived the idea and performed most of the large-scale genomic analyses. MR wrote the scripts for post-processing of the data, performed simulations and wrote the manuscript. VK performed all the PCR experiments and helped to develop genome analysis methods. FDP provided help with data management and visualization. All authors read and approved the final manuscript.

### Ethics approval and consent to participate

The genome data were collected and used with ethical approval (Nr. 206T4, obtained for the project SP1GVARENG).

### Consent for publication

Not applicable.

### Availability of data and materials

All scripts (pipeline_ref_plus.sh, pipeline_ref_minus.sh and pipeline_merging_and_filtering.sh) and software (gtester) created for this study are available from GitHub (https://github.com/bioinfo-ut/AluMine). The FastGT package used for genotyping the Alu insertions is also available from GitHub (https://github.com/bioinfo-ut/GenomeTester4/blob/master/README.FastGT.md). *K*-mer lists for genotyping Alu elements using FastGT are available from University of Tartu webpage (http://bioinfo.ut.ee/FastGT/). The whole genome sequencing data that support the findings of this study are available on request from Estonian Genome Centre (https://www.geenivaramu.ee/en) but restrictions apply to the availability of these data, and so are not publicly available.

### Additional Files

*Puurand_2019_AdditionalFile1.pdf*

Additional file 1. Figure S1 and Figure S2 explaining the REF-and REF+ discovery algorithms. (PDF, 139 kb)

*Puurand_2019_AdditionalFile2.xlsx*

Additional file 2. Supplementary tables Table S1, Table S2, Table S3 and Table S4. (XLSX, 2.1 Mb)

