## Additional File 1 for "AluMine: alignment-free method for the discovery of polymorphic Alu element insertions"

### Additional File 1. Supplementary Figures

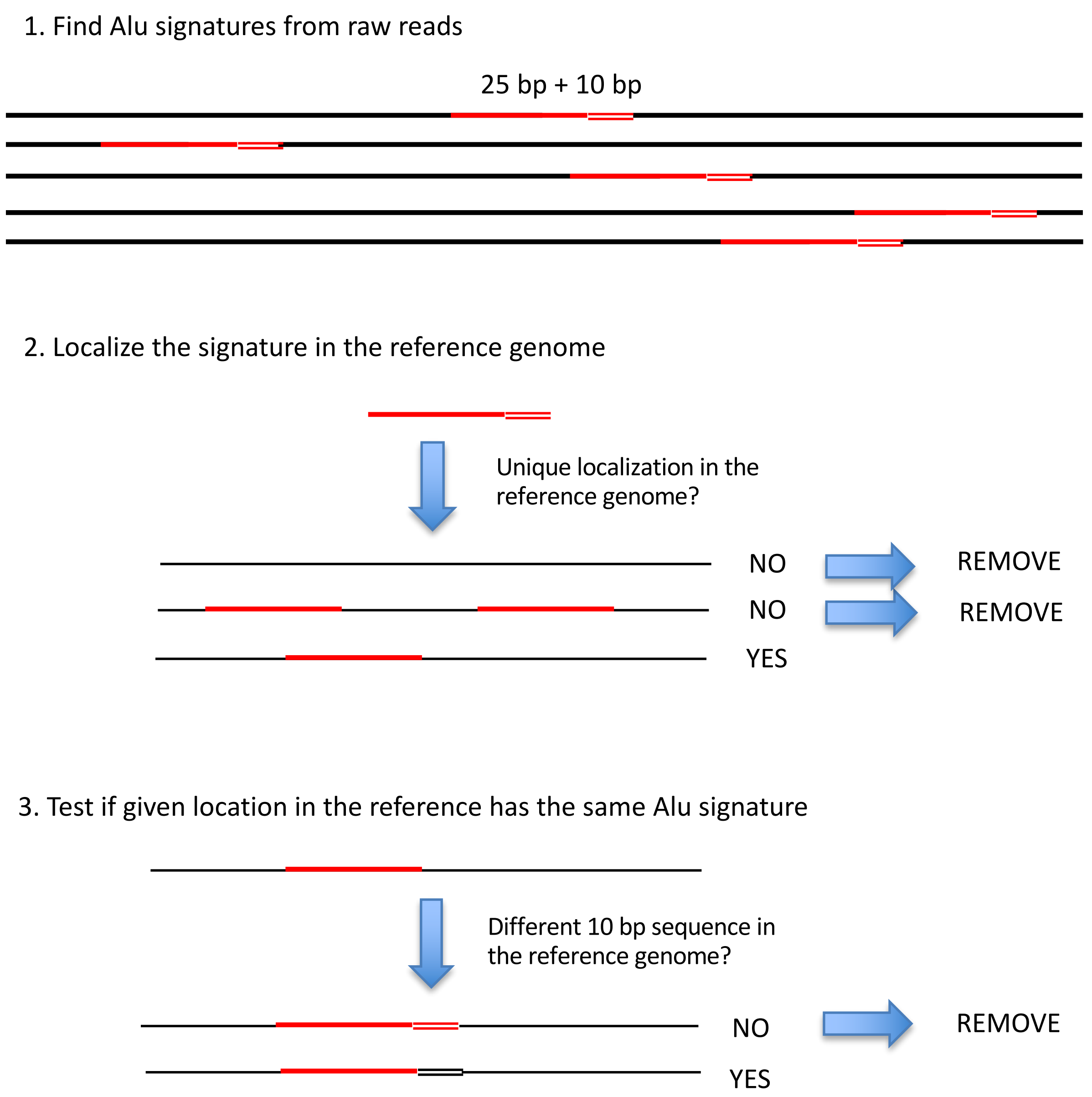


**Figure S1. Main steps of the REF– discovery pipeline.**

<u>Step 1.</u> Search for 10 bp Alu signature sequences in raw reads from sequenced individuals. Extract 25 bp sequence from the 5′-flanking region of the signature sequence and add it to the 10 bp signature. Remove the candidate if the frequency of the resulting 35-mer in a given individual is <5 or >100.
<u>Step 2.</u> Use the 25 bp region to determine the location of the Alu element in the reference genome using gtester4. Remove the candidate if its location is not detectable in the reference genome (Alu elements from heterochromatin). Remove the candidate if the 25 bp sequence is present in multiple locations in the reference genome (Alu elements from repeated regions).
<u>Step 3.</u> Extract the 10 bp sequence from the reference genome. Compare it with the Alu signature. Remove the candidate if the reference genome already contains an Alu element in this position (fixed Alu elements).

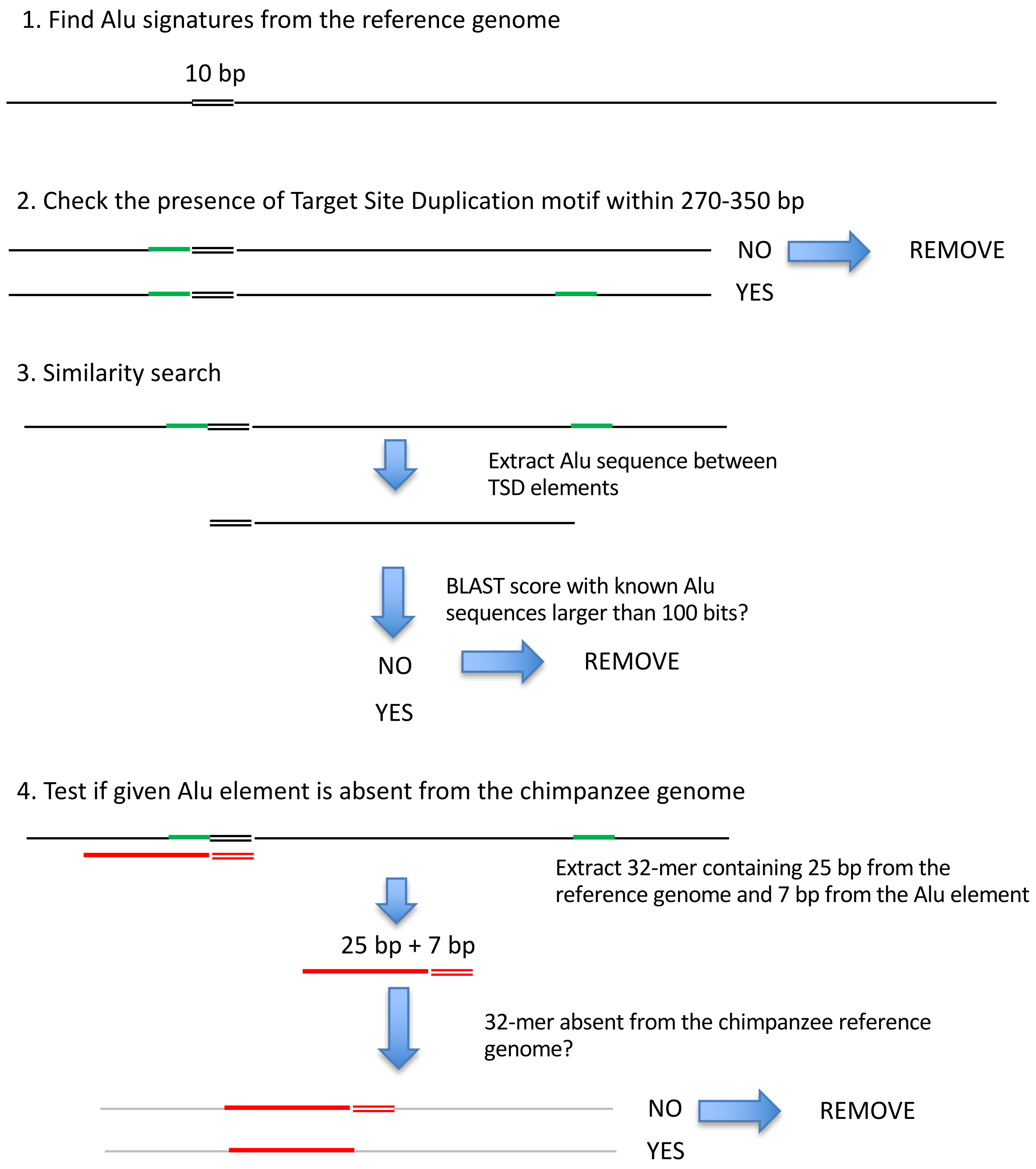


**Figure S2. Main steps of the REF+ discovery pipeline.**

<u>Step 1.</u> Find all 10 bp Alu signature sequences, allowing up to 1 mismatch with the reference genome.

<u>Step 2.</u> Identify the 5 bp target site duplication (TSD) sequence at the 5' end of the Alu signature sequence. Search for identical 5 bp TSD sequences at the 3' end of the Alu element. Remove the candidate if the 3' end TSD is not detected within 270 - 350 bp of the start of the Alu signature sequence.

<u>Step 3.</u> Test the similarity between detected Alu elements with known Alu elements.

<u>Step 4.</u> Generate REF+ *k*-mers (25 nt from the genome and 7 nt from the Alu sequence) for each candidate. Count the frequencies of these *k*-mers in the chimpanzee genome, allowing 2 mismatches. Remove the candidate if the 32-mer was detected at least once in the chimpanzee genome.
